# Molecular noise modulates transitions in the cell-fate differentiation landscape

**DOI:** 10.1101/2025.07.28.667061

**Authors:** Y. Liu, A. Zanca, M.P.H. Stumpf, L. Ham

## Abstract

Waddington’s epigenetic landscape has become one of biology’s cornerstone metaphors, widely used both conceptually and computationally. Cell types are often associated with the stable stationary points or valleys of this landscape. In previous work, we showed that the molecular noise dominating sub-cellular dynamics can distort and profoundly reshape this landscape. In non-equilibrium systems, an equally profound question arises: to what extent does noise alter the transition paths between valleys in such dynamic landscapes. We tackle this question using a set of illustrative exemplars, and show that noise gives rise to paths that differ substantially from the canonical least-action paths calculated under deterministic dynamics. We dissect the dynamics of these exemplars and determine the reactive density and transition currents, which show us, respectively, where and how transitions occur for different realisations of stochastic dynamics. Our analysis unambiguously demonstrates that reaction paths for stochastic dynamics diverge non-trivially from their deterministic least-action paths or simple barrier crossing models.

## I. INTRODUCTION

During early life and development, embryonic stem cells can differentiate into all the different cell types within the body [1]. Tissue-specific stem cells can generate all or at least subsets of the cells in a given tissue and are important during growth, tissue homeostasis, and tissue repair and regeneration [2]. The process by which an embryonic or tissue-specific stem cell ultimately gives rise to specific cell types is known as stem cell differentiation. This process is tightly regulated and involves a cascade of events that lead to the activation or repression of specific genes. These genes determine the characteristics of the specialized cell, such as its function, structure, physiological and spatial niche [3–5]. The genes and proteins that a cell express reflect, but also shape, all these biophysical, biochemical, and cell phsiological characteristics.

As a convenient shorthand, we often identify the corresponding cellular states (such as expression patterns) with the phenotype of a cell [6]. In the context of cell differentiation, for example, the cell’s phenotype is often assumed to be characterized by its gene expression profile, which is influenced by both intrinsic factors (such as the cell’s genetic makeup) and extrinsic factors (such as signals from neighbouring cells or the surrounding environment) [7, 8].

One of the central questions in developmental biology is to understand how the correct cell types appear at the right place and time in the correct numbers and proportions. To attain a given cell type, a cell (or lineage) must therefore traverse a particular differentiation path (out of a vast number of such potential paths) in a self-organised manner through activiation of the correct signalling and gene regulation networks [9]. Decoding this complex biological transition process can help us identify key regulatory mechanisms of cell differentiation. This has fundamental importance as well as transformative potential for applications in medicine, tissue regeneration and biotechnology. For example, in revealing how healthy cells transition into diseased states such as cancerous states [6, 10–12], or how to control transitions between different cellular states for tissue repair or organ regeneration [13–15].

The dynamics of cell differentiation [16] have been studied predominantly through the lens of (deterministic) dynamical systems theory. This perspective originates from Waddington who introduced the “epigenetic landscape” as a metaphor for cell differentiation. Since then, most modelling of cell fate has focused on “quantifying” the landscape where the valleys in the landscape correspond to stable cell states (differentiated cell types) and the barriers between valleys correspond to energy barriers that cells must overcome to transition between different states. Quantifying the potential landscape involves determining the “energy” (where “energy” is not physical energy, but used as an analogy representing differentiation potential) of different cell states and the barriers between them, which can provide insight into the stability of cell states [17–19].

The transition path for stem cell differentiation can be understood as the series of intermediate states or stages that a stem cell passes through as it changes from a less differentiated state to a more specialized state [20]. This path is often depicted by a trajectory in a suitable high-dimensional space representing the changing gene expression profiles of the cells throughout the course of differentiation [16]. In a dynamical system, a transition pathway can be understood as the route that a cell transits from one stable attracting state to another on the potential landscape, or another suitable manifold.

Cells are inherently noisy systems, with fluctuations arising from the stochastic production and degradation of molecules, as well as interactions with neighboring cells. This noise can profoundly reshape the structure of the epigenetic landscape, including in non-intuitive ways [21]. Traditionally, the landscape has been viewed as being shaped by deterministic forces, which are characterized by a quasi-potential. However, noise interacts with deterministic forces, changing the structure of the landscape. Such changes include creating or destroying attracting states and promoting or inhibiting transitions by changing the heights of barriers between attracting states.

Exactly how noise changes transition pathways is still not well understood [22, 23]. To study transitions between cellular states, the minimum action path (MAP) has been used under the assumption that a quasi-potential exists [24]. Motivated by the large deviation principle, the MAP measures the minimum energy that is required for a system to transition between attractor states [25, 26]. It has been used to identify the most probable pathways between different cell states, effectively providing a deterministic approximation of rare stochastic transitions [24, 27–30]. The advantage of this approach lies in its computational efficiency and adaptability to high-dimensional spaces. However, it has important limitations when analyzing the role of noise in transitions. Firstly, the MAP is only valid in the small noise limit. This assumption often does not hold in biological systems, where noise can be substantial and system-altering. Second, it offers only a coarse approximation by capturing the exponential scaling of transition rates, without resolving finer details of the dynamics (See Appendix A). Most importantly, the MAP describes a single, optimal path and thus fails to capture the full range of possible transition dynamics. As illustrated in Fig. 1 multiple transition paths may contribute meaningfully to differentiation. The MAP alone cannot reveal the distribution or relative importance of alternate transition pathways, limiting its ability to describe the diversity and richness of real stochastic transitions.

**FIG. 1.**
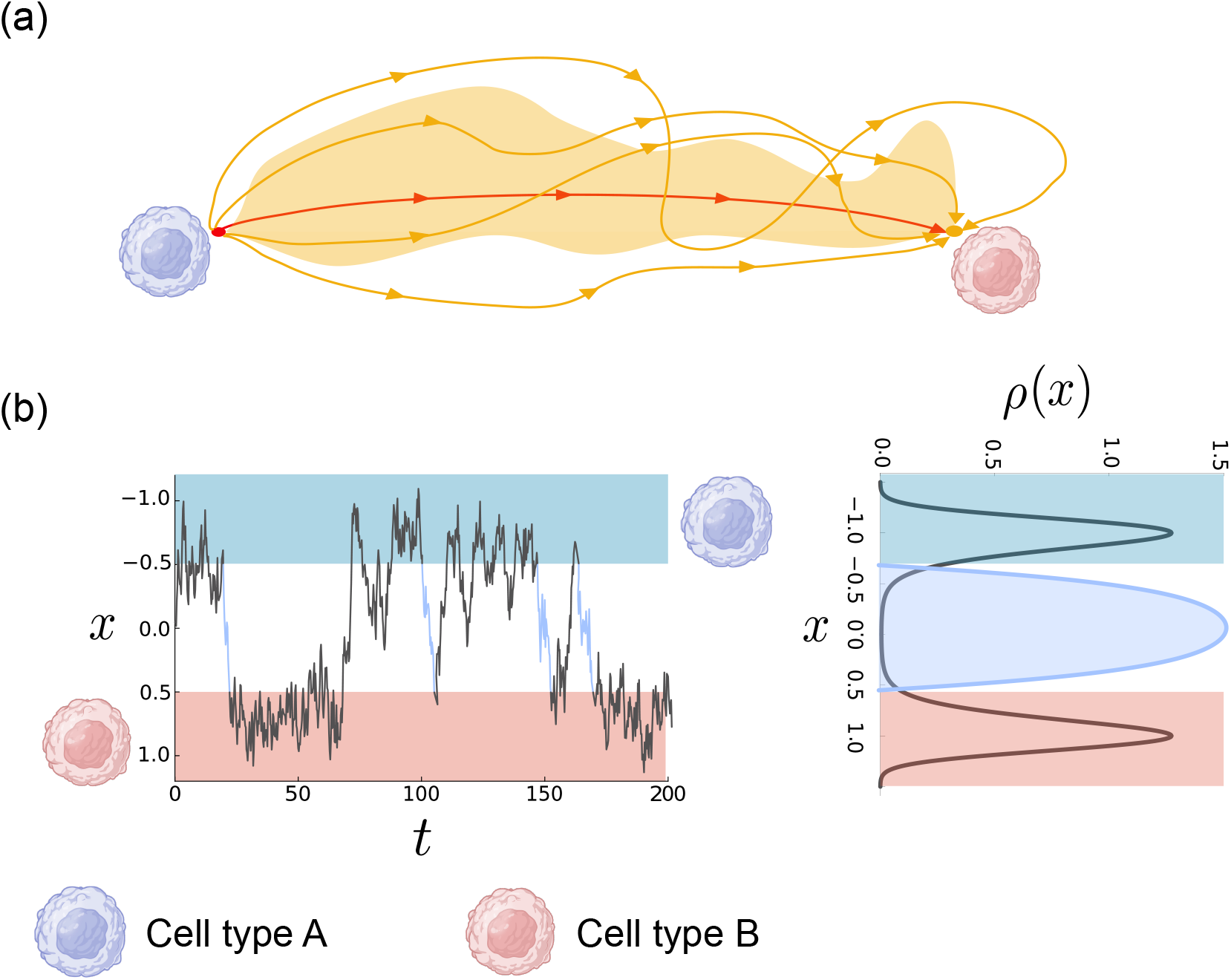
Example trajectories of transitions for the cellular dynamics in a double-well potential system. (a) Transition trajectories begin at the initial state *A* indicated by the blue cell, transition towards the target state *B*, indicated by the red cell. Yellow paths represent example transition trajectories, while the red path represents the MAP of transitions. The yellow shaded area represents the distribution of transition pathways or reactive density in TPT. (b) Left panel: Simulated trajectories with light blue areas representing transition pathways. Blue and red shaded regions represent the cellular states *A* and *B*, respectively. Right panel: The stationary distribution of a cell-fate process (black curve) and the distribution of the transition pathways for simulated trajectories (light blue shaded curve).

Here, we employ transition path theory (TPT), which characterizes the statistical properties of transitions between two attracting states. TPT allows us to consider the full ensemble of transition paths connecting two attracting states. TPT has been successfully applied in many biological systems, including extracting transition pathways in protein folding, identifying parallel transition routes in epithelial-to-mesenchymal transition, and quantifing the likelihood of all possible transition trajectories between cellular states using single-cell data [31–33]. TPT was originally developed as a purely theoretical perspective. The introduction of RNA velocity enables the reconstruction of cellular landscapes from scRNA-seq data by deriving vector fields of cellular dynamics [34]. Recent studies have used TPT to characterize transition dynamics in single-cell datasets, revealing key insights into the reprogramming of gene expression. In particular, TPT has shown that interactions between cell communities intensify during cell-fate transitions, coordinating global gene expression changes and guiding cells toward distinct developmental trajectories [35].

Despite the broad applicability of TPT, few studies have examined how noise influences transitions within this framework. In this work, we systematically map the impact of noise on transition pathways using TPT. Our findings show that noise does more than modulate barrier heights between attractor states, it can fundamentally re-shape the transition landscape, revealing alternative intermediate states and pathways that differ markedly from those observed in the absence of noise.

## II. FROM CELL STATES TO TRANSITION PATHWAYS

### A. Waddington’s landscape

C.H. Waddington introduced the epigenetic landscape as an intuitive way to reason about cell differentiation [16]. In this metaphor, the landscape consists of continuous hills and valleys, where each point in the landscape represents a cellular state. Cells undergoing differentiation can be conceptualized as marbles rolling down a hill from pluripotent states (cells that can give rise to all cell types) towards a stable, differentiated state in the valleys. The trajectory a cell takes represents its differentiation pathway. Along any path, barriers (higher ridges) confine cellular behaviour. These barriers can reflect biological controls that constrain cell types. Gene regulation, signalling, and effects from the microenvironment guide cells through the differentiation process, and will render the landscape/manifold as dynamic rather than static [23].

Inspired by Waddington’s metaphor, dynamical systems theory has been employed to describe the developmental process of a cell. Specifically, gene regulatory networks (GRNs) determine the landscape structure by assuming that cell fate is governed by these networks. The GRNs, in turn, can be expressed mathematically by systems of differential equations. The behaviour of dynamical systems with noise can naturally be described by a stochastic differential equation (SDE) with parameters *θ*:

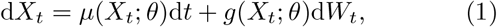

where *X*_*t*_ represents the state of the system (for GRNs, typically defined in terms of molecular concentrations) at time *t, µ*(*X*_*t*_; *θ*) represents the deterministic forces governed by the GRNs and *g*(*X*_*t*_; *θ*) is the diffusion term accounting for the intrinsic stochasticity. The term *dW*_*t*_ refers to increment of a Wiener process which satisfies *E*[*dW*_*t*_] = 0 and *E*[*dW*_*t*_*dW*_*s*_] = *δ*(*t* − *s*).

From the perspective of the potential landscape the dynamics are often modelled as gradient systems with an associated potential function:

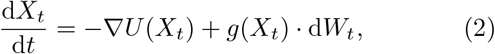

where the potential function *U* (*X*_*t*_) characterizes the deterministic evolution of cell states and identifies the final attracting states the system tends to evolve towards. Note that we drop *θ* from the notation for brevity.

The parameters, *θ*, can substantially influence the dynamics of networks by altering the positions of attractors (stable states) and saddle points (decision points). These parameters can be regarded, for example, as activation or inhibition factors within GRNs, which helps define the structure of the landscape. The transitions from one cell state to another can be seen as switches between stable states in the underlying dynamical system, effectively driven by intrinsic noise in the system.

For gradient systems, the stationary (invariant) distribution can often be expressed as the Gibbs-like distribution[16, 36],

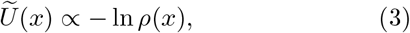

where *ρ*(*x*) is the stationary distribution and 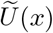 is the potential function of the landscape under the influence of noise.

For diffusion in one dimension, the stationary distribution can be expressed as [21]:

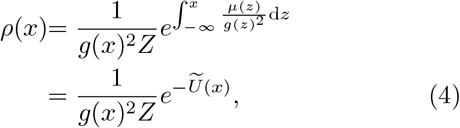

with

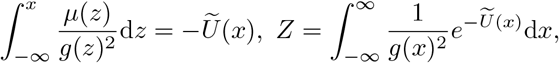

where *Z* is the normalization constant.

Therefore, one can approximate the quasi-potential landscape 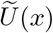 from the stationary distribution *ρ*(*x*) by conducting simulations of specific models or collecting experimental data and performing density estimation via e.g. Eqn. (3) [37] (note that in one dimension every dynamical system is necessarily a gradient system).

While intrinsic noise during cell differentiation is incorporated in SDEs of the form given in Eqn. (1), most existing studies assume additive noise (i.e. *g*(*X*_*t*_; *θ*) = *σ*), even for biological systems where we know that this is unfounded [22]. Biological noise is not constant but can vary depending on cellular states, gene expression levels, and gene interactions, among other factors. To obtain an accurate description of cell fate, it is necessary to incorporate multiplicative noise, which captures state-dependent properties, into our models.

This raises a key question: how does multiplicative noise affect the cell-fate differentiation landscape? A recent study has shown that different types of noise can influence the landscape in distinct ways. While additive noise can only quantitatively change the structure of the landscape (i.e. valleys become shallower and hills become flatter), multiplicative noise can qualitatively reshape and distort the landscape [21]. Additive noise will not change the basins of attraction of a landscape, whereas multiplicative noise can move the attracting states to different locations, induce stochastic bifurcations that divide attracting states into multiple stable states, or merge existing attracting states into a single new state.

Previous analyses, however, provide only a static view of cellular states, without addressing the dynamics of transitions between them. Key questions remain unresolved such as how long it takes for a cell to transition between states, which pathways are most likely, and where cells tend to dwell along these trajectories in the presence of noise. As transitions between basins of attraction are rare events [36], directly capturing them through simulation is highly challenging. To address this, we turn to TPT, which provides a principled framework for analyzing noise-driven transitions between cellular states.

### B. Transition path theory

TPT was introduced to study the reactions in molecular, including biological, systems where transitions are rare. By solving the backward and forward Kolmogorov equations, TPT captures those regions in state space through which transitions are more likely to pass, and regions in state space where the systems spends most of the time. Transition properties are evaluated in terms of a forward committor function *q*^+^(*x*), which is the probability that a cell starts in state *A* and reaches state *B* before returning to state *A*. We use Ω to denote the state space and *∂A, ∂B* to denote the boundaries of states *A* and *B*. The forward committor function can be written as

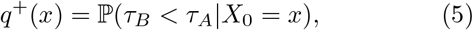

where *τ*_*A*_ = inf {*t* ∈ ℝ_+_ |*X*_*t*_ ∈*∂A*} is the first hitting time of a diffusion process (*X*_*t*_)_*t* ≥0_ that starts at a point *x* ∈ Ω_*AB*_ = Ω\(*A B*) and similarly for *τ*_*B*_.

One can show that *q*^+^(*x*) is the solution of the Dirichlet problem [38, 39] with

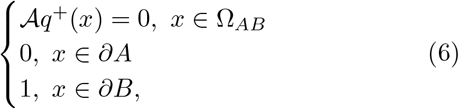

where 𝒜 is the generator of the diffusion process defined as

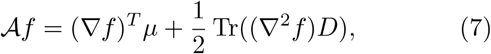

where *µ* is the deterministic force and *D* is the diffusion matrix of the process. We use the finite element method to numerically mesh the space to solve Eq. (7) and approximate the forward committor function.

Similarly, the backward committor function *q*^−^(*x*), which can be solved by the time-reversed generator, is defined as the probability that the diffusion process arriving at a point *x* originates from within state *A*. If the process is time-reversible then we have *q*^+^(*x*) = 1−*q*^−^(*x*).

By analyzing the statistical properties of the committor functions – including reactive density, transition currents, transition rate, and mean first passage time – we can draw conclusions about the transition paths between cell states.

We say that the diffusion process (*X*_*t*_)_*t* ≥0_ is in transition from state *A* to state *B* if it was previously in state *A* and is transitioning towards state *B* before it is possible to return to state *A*. The probability density function of such transition processes can be defined as:

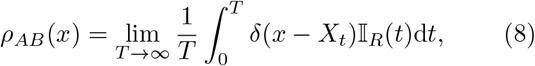

where the indicator function 𝕀_*R*_(*t*) is 1 if it is in transition and 0 otherwise.

By definition *ρ*_*AB*_(*x*) is exactly the probability that at point *x, X*_*t*_ will visit *B* first before *A* and that *X*_*t*_ comes from *A*, restricted by the probability density function of the whole system at point *x*. Therefore, the probability distribution of the transition pathways is [40]:

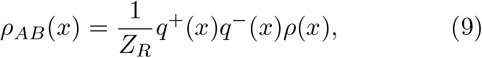

where *Z*_*R*_ = ∫*q*^+^(*x*)*q*^−^(*x*)*ρ*(*x*)d*x* is the normalization constant that defines the probability that the process is found in the transition from *A* to *B* at any random time point. Note that we refer to *ρ*_*AB*_(*x*) as the *reactive density* below to distinguish it from the stationary density function *ρ*(*x*) and to make the termininology consistent with existing literature [38–40]. The reactive density tells us where in state space cells spend time during the transition as shown in Fig. 1.

The probability flow associated with the transition process can be studied via the probability current, *J*_*AB*_, determined by the transition path process as [40, 41],

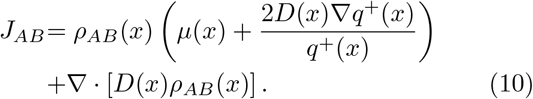

Under potential theory, the stationary distribution and probability current define the intrinsic structure of the potential landscape; we refer to this as the *transition current*. Cellular states with a higher value of density *ρ* can be understood as the attracting states in the landscape. A higher transition current at point *x* can be understood as a higher flow of probability through a point *x*. Therefore, cellular states with higher transition currents are less stable and more likely to be shortlived. The transition current therefore tells us how transitions between cell states occur.

Regions with higher reactive density can be understood as transition states in the landscape. Transition states with low transition probability current can be viewed as transition traps of the process, as it is more difficult to transition out of these states.

Since we are interested in the effect of noise on the expected time of transitions, we define

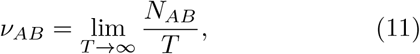

as the transition rate which quantifies the mean number of transitions until time *T*, where *N*_*AB*_ denotes the total number of transitions between attracting states in [0, *T*].

We also define

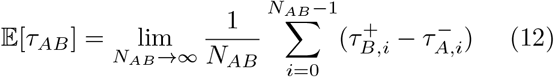

as the mean first passage time (mfpt) from *A* to *B* where 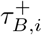 is the *i*^*th*^ entrance time to *B* and 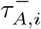 is the *i*^*th*^ exit time from *A*. We then have

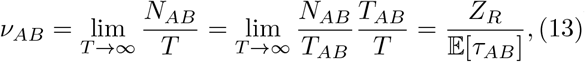

where *T*_*AB*_ is the total time that a trajectory is in transition from *A* to *B*. The normalization constant *Z*_*R*_, defined previously, can be interpreted as

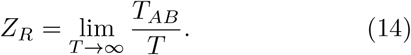

It follows from the definition of the current traversing any surface *S* separating *A* and *B*, that the transition rate *ν*_*AB*_ is the total flux through the surface i.e. [41, 42]

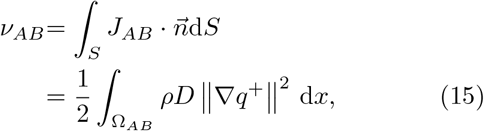

where 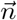 is the unit normal pointing outwards from the surface *S*.

To study transitions between cellular states in cell differentiation under different types of noise, we will use TPT and quantify the statistical properties of the transition pathways described above. We use Eqn. (9) to quantify the distribution of transition pathways and Eqn. (10) to quantify flows. Additionally, Eqn. (12) and Eqn. (15) will be used to assess transition times under different noise conditions.

## III. RESULTS

### A. Gradient systems in equilibrium

We start by considering the system that is in equilibrium (see Appendix B 1) with canonical one-dimensional double-well potential

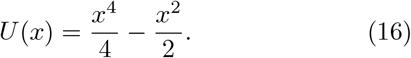

This system has two distinct local minima at *x* = ±1, separated by a saddle point at *x* = 0. Therefore, the transition from one state to another can be understood as moving the state space variable *x* from one minimum to the other. For simplicity we define the neighbourhood around the minimum *x* = −1 as state *A* and *x* = 1 as state *B* and consider transitions from *A* to *B*.

In one dimension we can show that

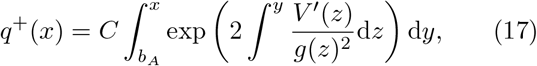

solves Eqn. (6), where *b*_*A*_ is the upper bound of the basin of attraction of state *A* and *C* ∈ ℝ^+^ is the normalization constant that enforces *q*^+^(*x*) ∈ [0, 1], ∀_*x*_ ∈ Ω_*AB*_.

The equilibrium property of the system gives the backward committor function as

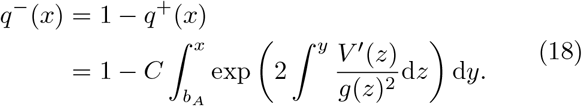

Therefore, the reactive density *ρ*_*AB*_(*x*) can be obtained from Eqn. (9) via numerical integration.

#### 1. Transition promotion with increasing noise strength

We first consider the case of additive noise, *g*(*x*) = *σ*, with noise strength *σ >* 0. The reactive density, *ρ*_*AB*_(*x*), given in Fig. 2(a), displays the transition dynamics for increasing values of *σ*. We can see that *ρ*_*AB*_(*x*) has a peak at the saddle point, *x* = 0, indicating that the region around the saddle point serves as the transition state for this system. When the noise strength *σ* is low, the density is heavily concentrated at the saddle point with wings extending to its surroundings. That is to say, the system will be trapped at the transition state in the absence of noise where reactive density acts like a Dirac delta function centered at the saddle point. The transition can only take place when there is some noise associated with the system. Noise breaks the stability of the transition state at the saddle point, allowing the cell to transition between states *A* and *B*. A transient change in the density peak around the saddle point is observed with increasing noise strength, as shown in Fig. 2(a) (left panel). This suggests that the stability of the transition state decreases with increasing noise strength initially, but subsequently begins to increase and stabilize. The stability of the transition state can be quantified using the value of the reactive density at the saddle point, shown in Fig. 2(c). Our results reveal that the transition is maximally promoted when the noise strength is *σ* ≈ 0.43 and thereafter the stability of the transition state will gradually increase towards an asymptotic limit as the noise strength grows, corresponding to an increase in the reactive density.

The In the case of multiplicative noise, we consider 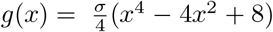, again with noise strength *σ >* 0. The introduction of multiplicative noise into the system can lead to a qualitative change in the structure of the landscape [21]. Specifically, here the positions of the attracting states are altered: the minima in the original landscape *ρ*(*x*) are pushed away from *x* = ±1 towards 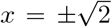 as noise strength increases. We consider the shifted minima as new attracting states and label them as states *A* and *B*, as before.

**FIG. 2.**
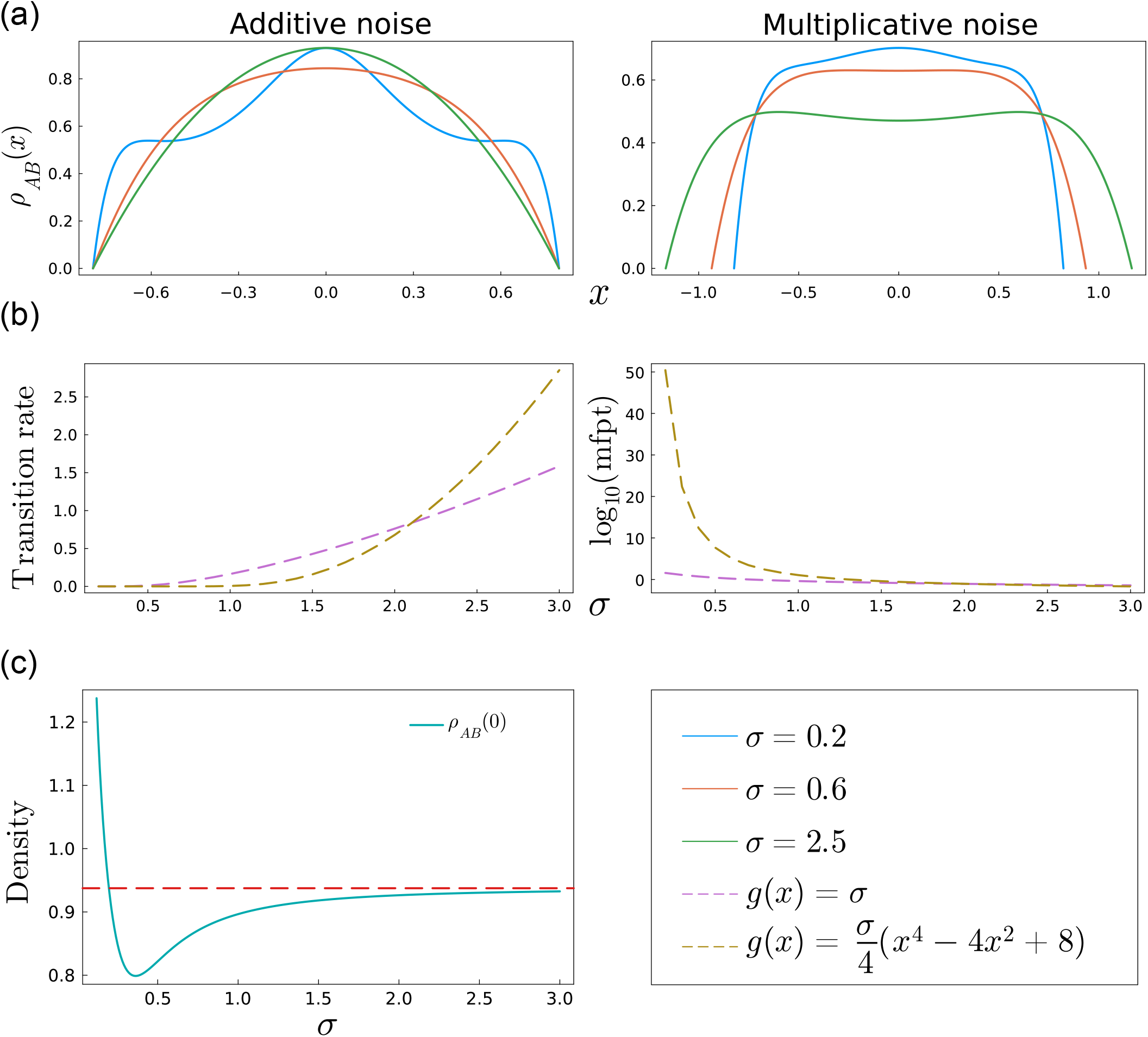
Reactive density, transition rate and stability analysis for the one-dimensional double-well potential with additive and multiplicative noise. (a) The reactive density, *ρ*_*AB*_, (*x*) of the diffusion process under different types of noise. Left panel: additive noise with *g*(*x*) = *σ*. Right panel: multiplicative noise with 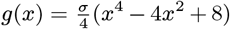. As noise strength *σ* increases, the maximum of *ρ*_*AB*_ (*x*) concentrates around the saddle point *x* = 0 for additive noise. While for multiplicative noise, the maximum is bifurcating towards two maxima around the saddle point. (b) The transition rate, *ν*_*AB*_, and mean first passage time, *E*[*τ*_*AB*_], in log_10_ scale for different form of noise. With the increase of noise strength *σ*, the transition rate is increasing monotonically and the mean first passage time is decreasing monotonically. In the small noise limit, the system with multiplicative noise is less likely to have transitions than a system with additive noise. Conversely, for the large noise limit, the multiplicative noise system is more likely to have transitions. (c) The reactive density for additive noise at the saddle point (transition state) under different noise strength *σ*. The red dashed line represents the asymptote for large noise strength. The minimum value of the reactive density, correpsonding to lowest stability, is attained at *σ* ≈ 0.43.

Figure 2(a) (right panel) gives the reactive density of the multiplicative noise system under varying noise strength. Similarly to the case of additive noise, with low noise strength, the transition state is again concentrated around the saddle point. However, rather than being tightly localized around the saddle point, the transition state region expands with increasing noise strength, covering a broader neighbourhood around the saddle point due to the interplay between determinisic forces and multiplicative noise. Moreover, unlike the case of additive noise, where the stability of the transition state exhibits transient changes with increasing noise strength, the stability of transition states continuously decreases with increasing noise strength.

The destabilization of the transition state leads to a bifurcation where the original transition state is split into two states symmetrically arranged around the original saddle point at *x* = 0. Thus under high enough multiplicative noise, there are two transition states located at points equidistant from the original saddle point, which is no longer a transition state.

Analyzing the transition rate, *ν*_*AB*_, and mean first passage time, *E*[*τ*_*AB*_], for both additive noise and multiplicative noise (Fig. 2(b)) reveals that increasing noise strength, *σ*, generally promotes transitions. This finding agrees with a previous study [21], which showed that larger noise strength can lower the transition barrier, therefore promoting transitions. Additionally, we find that when the noise strength is low, transitions are less likely to take place under multiplicate noise than additive noise. However, when noise strength is large enough, transitions occur more frequently with multiplicative noise. This behaviour further suggests that, while there are more transition states due to bifurcation under multiplicative noise, these transition states are less stable than the original one at the saddle point and therefore become more permissible to transitions between states.

We now formalize this intuition about how the structure of the reactive probability density changes under multiplicative noise (see Appendix C for further details). Specifically, we have

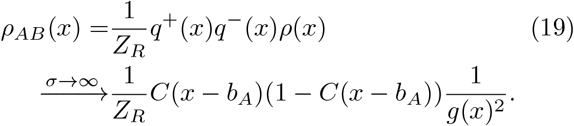

The mode of the reactive probability density function will approach the mode of

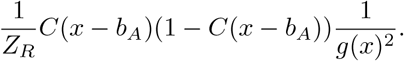

### B. Gradient system in non-equilibrium

We now extend our study to higher-dimensional systems where multiple transition pathways may exist, subject to the direction of probability flux (see Appendix B 2). Here, we approximate the numerical solution of Eqn. (6) using the finite element method (see Appendix D).

We consider the two-dimensional double-well potential given by:

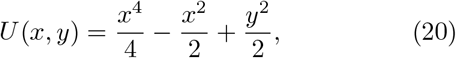

which consists of a double-well potential in the *x*-direction and a parabolic function in the *y*-direction. Due to the higher-order terms in the *x*-direction, the potential function is primarily influenced by variations along *x*-direction. The landscape again has two minima at (±1, 0) that are separated by a saddle point at the origin, (0, 0).

The attracting states, *A* and *B*, are defined as the basins around the two minima of the probabilistic landscape at (±1, 0) respectively. Our analysis is conducted over the domain Ω = [−2.5, 2.5] × [−2.5, 2.5], ensuring sufficiently high potential values at the boundaries. This results in the stationary probability density function *ρ*(*x, y*) effectively vanishing around the boundaries of the domain.

#### 1. Unstable saddle point with increasing noise

We start with the simpler case where only additive noise is present in the system for both the *x*- and *y*-directions

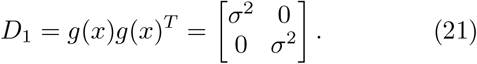

In this case, the structure of the landscape remains unchanged with states *A* and *B* at (±1, 0) separated by the saddle at (0, 0) (Fig. 3(a), left panel).

**FIG. 3.**
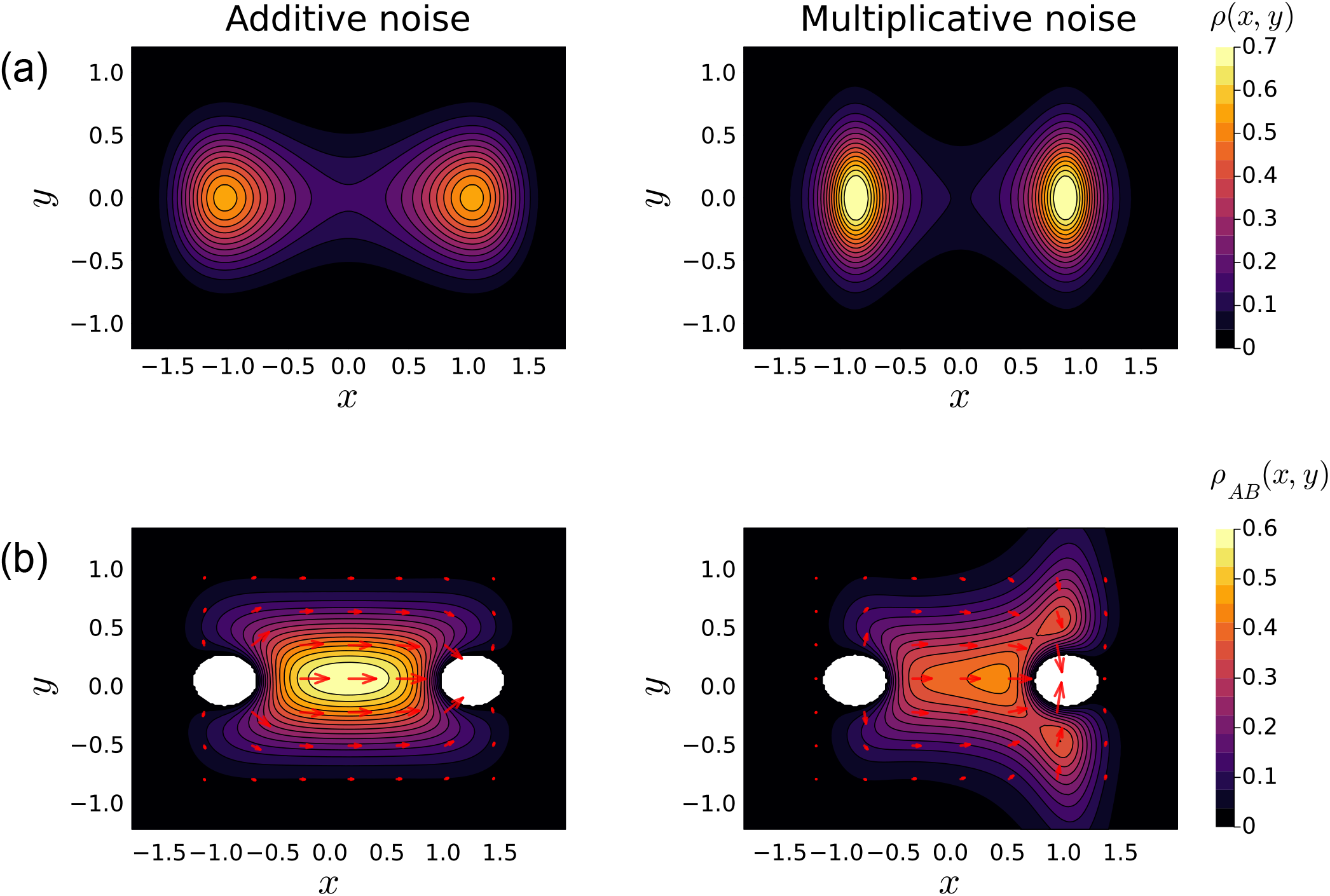
The stationary probability density function *ρ*(*x, y*) and the reactive density *ρ*_*AB*_ (*x, y*) for the two-dimensional double-well potential system with different noise terms. (a) The stationary probability density function with additive noise described by diffusion matrix *D*_1_ on the left and multiplicative noise described by *D*_2_ on the right. (b) The reactive density *ρ*_*AB*_ (*x, y*) for with additive noise on the left and multiplicative noise on the right. The white regions correspond to the attracting states *A* and *B* respectively. The red arrows represent the probability current *J*_*AB*_ of the transition processes. The larger the arrows, the higher the probability that cells will transition through these regions. Introdution of multiplicative noise into the system leads to the creation of new transition states symmetric around the target attracting state as well as enabling new transition pathways towards the target attracting state. The noise strength *σ* is set to be 0.6 here for both additive noise and multiplicative noise.

Note that when the additive noise is coupled with the gradient drift, the system is in equilibrium [43] and conseqently the backward committor function can be simplified to

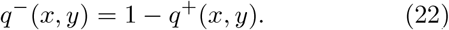

Figure 3(b) (left panel) gives the contour plot of reactive density *ρ*_*AB*_(*x, y*) alongside the transition probability current *J*_*AB*_, represented by red arrows where the value of the flux is indicated by the arrow size. The larger the arrows, the higher the likelihood that cells will transition through these regions.

One can see that the transition pathways in this system are mostly through the central region crossing the saddle point which directly connects states *A* and *B*. This is represented by both the higher reactive density *ρ*_*AB*_(*x*) around the central region and the higher probability current *J*_*AB*_ crossing this region.

Therefore, the area around the saddle point can be viewed as the transition state of this system. Additive noise does not change the transition dynamics or the topology of the reactive density function, similar to the previous section. Its influence on the transition dynamics is mainly in reducing the stability of the transition state with increasing noise strength, making it easier for a cell to transition from state *A* to *B*. This is represented by both a decrease in reactive density and an increase of flux through the transition state (Appendix E).

#### 2. Creation of new transition states and pathways

Now we turn to the case of multiplicative noise where we take

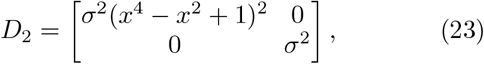

as the noise in the system. While the intrinsic features of the landscape are maintained – the system again has two attracting states separated by a saddle point in between – multiplicative noise alters the position of the attracting states where minima are pushed away from (±1, 0) towards 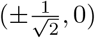 with increasing noise strength.

Although the overall intrinsic appearance of the landscape is preserved, multiplicative noise now also introduces new transition pathways as well as new transition states. The change in transition pathways is illustrated in Fig. 3(b) (right panel). When additive noise is present, the system exhibit a dominant transition pathway that connects attracting states *A* and *B* through the saddle point. However, when multiplicative noise is present, the transitions into state *B* will not proceed directly through the original transition state. Instead, the original pathways is bifurcated into two distinct and equally weighted branches, that vary in the y-direction, towards the target state. The two bifurcated pathways have equally weighted probability due to the symmetric multiplicative noise and potential. This result highlights the role of multiplicative noise in changing the transition dynamics of the system: it can create other transition routes and enable entrance into the desired attracting states through potentially different molecular spaces.

The creation of new transition paths can be understood as resulting from the interplay between noise and the state-dependent dynamics of the system. Multiplicative noise introduces state-dependent fluctuations, which lead to a change in (“effective”) curvature of the local landscape as well as the local stability. As a result, the interaction between the drift force and stochastic perturbations suggests that it may be worthwhile exploring multiple potential pathways, rather than concentrating on the highest probability one.

These findings have implications for understanding biological processes where noise is an intrinsic feature that depends on the cellular state. Previous studies [17] assumed transitions take place following routes mostly controlled by deterministic forces. Here we highlight the influence of noise on shaping the transition dynamics; this includes external or environmental sources of noise as new transition pathways can also be formed when cells are placed in a noisy environment. Therefore different noise environments can, in principle, lead to different transition states and transition pathways during cell differentiation.

### C. Reaction networks in equilibrium (Schlögl model)

So far, our study has focused on theoretical potential models where multiplicative noise *g*(*x*) is independent of the drift term, *f* (*x*). We now extend our analysis to biological systems where noise can appear as intrinsic or extrinsic. Regardless of the specific noise type, the system in one dimension remains in equilibrium.

To demonstrate this, we study the bistable chemical reaction system introduced by Schlögl

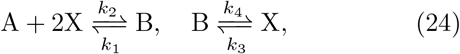

where *A* and *B* are chemical species with constant concentrations *a* and *b*, respectively, and *x* is the concentration of the species *X*, which is allowed to change over time.

This reaction system can be expressed as a gradient flow of a potential energy:

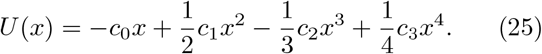

where the parameters *c*_3_ = *k*_2_, *c*_2_ = (*a* +3)*k*_1_, *c*_1_ = (*ak*_1_ + 2*k*_2_ + *k*_4_), *c*_0_ = *bk*_3_ are chosen such that the deterministic system exhibits two stable states, allowing for transitions between them.

When additive noise is incorporated into the system, the dynamics can be described by,

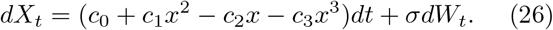

For intrinsic multiplicative noise, we consider the chemical Langevin equation (CLE) given in Eqn. (B8):

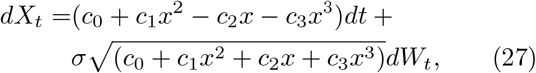

where the noise term *g*(*x*) now depends on the deterministic drift *f* (*x*).

#### 1. Sheared stability and transition states

Previous studies [21] have shown how different noise types can change the structure of the landscape for the Schlögl model. For purely additive noise the overall structure of the landscape is preserved but scaled, decreasing the barrier of transitions between two attracting states. Multiplicative noise causes a gradual change in landscape structure: the bistable structure of the landscape remains at low noise strength but becomes unimodal for large noise. This occurs because noise is significantly higher at the far right of the landscape compared to its strength around the origin which strongly pushing cells into a single region near the origin.

Regarding the transition dynamics, as shown in Fig. 4, at low noise strength, the system displays bimodal transition states for both additive and multiplicative noise, meaning that there are two distinct transition states associated with the system. However, these two transition states are not equally stable: the left transition state is more stable and corresponds therefore to higher probability density. This imbalance is even more evident with multiplicative noise which favours the transition state on the left.

**FIG. 4.**
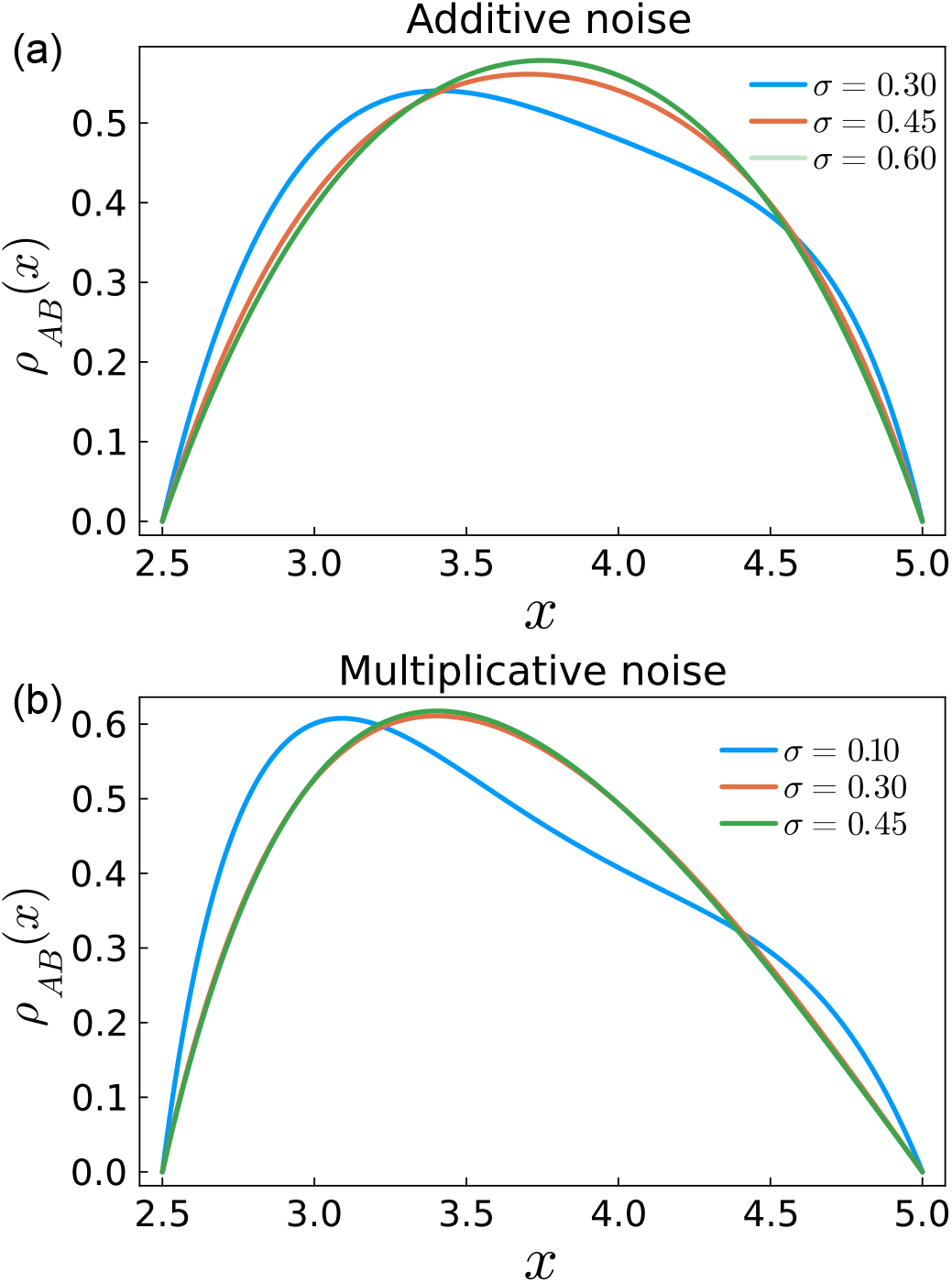
The reactive density *ρ*_*AB*_ (*x*) of the diffusion process of the Schlölgl model under different types of noise. (a) Additive noise with *g*(*x*) = *σ*. (b) Multiplicative noise with 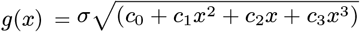. As *σ* increases, the maximum of *ρ*_*AB*_ (*x*) is concentrating around the saddle point *x* = 0 for additive noise while for multiplicative noise, the maximum is sheared and tends to be around the left. The parameters for the system are chosen as *c*_0_ = 0.4, *c*_1_ = 0.55, *c*_2_ = 0.20, *c*_3_ = 0.02. This results in a potential landscape with two attracting states for both additive noise and multiplicative noise.

At higher noise strengths, the two transition states will merge into a single state for both types of noise. For additive noise, this merged state forms a paraboloic function centered at the midpoint, consistent with the theoretical model in equilibrium as given in Section III A. For multiplicative noise, the merged state takes a sheared shape, skewing the stability of the transition state towards state *A* on the left. This result is similar to previous studies: the higher noise at the right extremes pushed the transition states towards the left.

Since the system is one-dimensional and in equilibrium, the limiting behavior of the reactive density function with high noise strength can be described by Eqn. (19). In this case, the resulting shape of the transition state comes from the interplay between the parabolic potential and the functional form of the multiplicative noise, highlighting the implicit influence of intrinsic multiplicative noise in shaping transition dynamics.

### D. Reaction networks in non-equilibrium (toggle switch model)

To further explore the effect of intrinsic noise on transition pathways in non-equilibirum chemical systems, we extend our analysis to a two-dimensional system. We examine the gene toggle switch model, which is used to describe binary cell-fate decisions. The system contains two genes, *X* and *Y*, that are self-activating while mutually inhibiting each other (Fig. 5(a)).

**FIG. 5.**
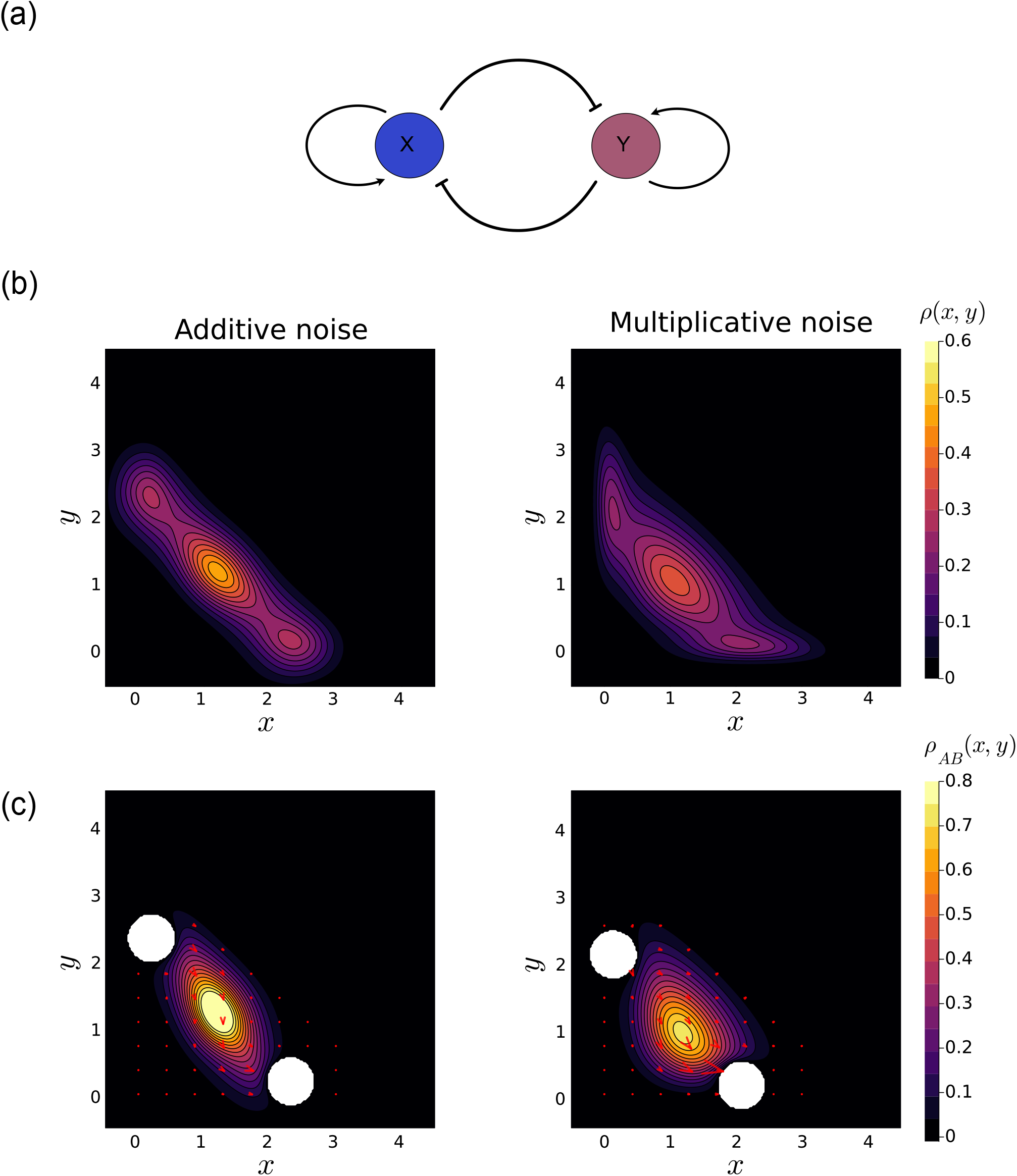
The stationary probability density and reactive density for the toggle switch model. (a) A schematic of the toggle switch model, consisting of two mutually repressing genes *X* and *Y* that also self activate. (b) The stationary probability density function *ρ*(*x, y*) for the toggle switch model with additive noise (left) and multiplicative noise described by the CLE (right). (c) The reactive density *ρ*_*AB*_ (*x, y*) for the toggle switch model with additive noise (left) and multiplicative noise (right). The white regions where arrows point outward and inward correspond to the attracting states *A* and *B* respectively. The red arrows with magnitude represents the probability current *J*_*AB*_ of the transition processes. The larger the arrows, the higher the probability that cells will transition through these regions. The introduction of multiplicative noise into the system leads to change in dominant pathway, from a winding movement into a heavily dominant one through the lower area of the transition state. The parameters for the system are *a* = 0.65, *b* = 1.2, *k* = 0.8, *s* = 0.6, *n* = 4, *g*_0_ = 0.2. The chosen parameters result in a landscape with three attracting states for both additive noise and multiplicative noise. The noise strength *σ* is set to be 0.45 here for both additive noise and multiplicative noise.

The dynamics of the system are described by the following SDE, where a Hill function is used to model the self-activation and mutual inhibition,

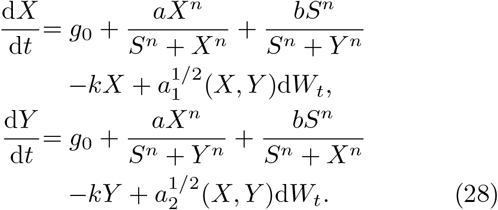

Here *g*_0_ is the basal synthesis rate for genes *X* and *Y, a* is their self-activation rate, *b* is their mutual inhibition rate, *k* is their self degradation rate, *n* is the number of regulators binding to the genes and *S* represents the threshold of the Hill function [44]; this is assumed to be the same for both genes.

The system is again considered with additive noise and multiplicative noise. For additive noise, the diffusion matrix is given by 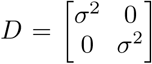. For multiplicative noise, the diffusion functions are defined through the CLE (see Appendix B 3), with:

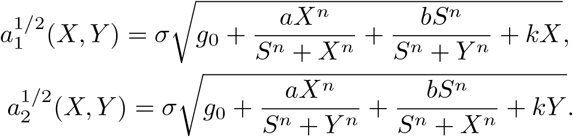

In both cases the system has three metastable states: two attracting states *A* and *B*, and an unstable transition state at the centre (Fig. 5(b)). We study the transition dynamics from state *A* to state *B*, through the central transition state.

#### 1. Change in dominant pathways

In the absence of noise, the MAP connects the two metastable states through the central unstable state; this follows the steepest descent path dictated by the potential gradient. When additive noise is introduced into the system, the overall structure of the landscape will be maintained. However, the curl force in the non-equilibrium system distorts the MAP from the steepest descent path and leads to a shifted pathway. Instead of following the steepest descent path, the probability current exhibits winding movements as it transitions through the transition state (Fig. 5(c) left panel). This observation under non-equilibrum conditions agrees with Wang *et al*.’s results [17], demonstrating that the MAP deviates profoundly from the steepest descent path under the influence of the curl force in non-equilibrium systems.

With the introduction of multiplicative noise formulated by the CLE to the system, the structure of the landscacpe is subtly altered from the original one while the intrinsic tristable structure is preserved: the three metastable states remain intact, without any bifurcations, destructions or creations of the attracting states. However, the dynamics of transitions is heavily reashaped by the intrinsic multiplicative noise (Fig. 5(c)).

With additive noise, transitions between attracting states follow a curved, undulating path, with relatively higher probability current coming out of state *A* and entering state *B*, while the current through the central transition state is reduced. This indicates that the transition state is relatively stable and transitions are less likely to occur when cells reach this transition state. By contrast, the probability current through the centre of the transition state increases under intrinsic multiplicative noise. The higher currents through the transition state, compared with those coming from state *A*, suggests easier transitions through the transition state and highlighting the decrease of stability at the transition state under intrinsic multiplicative noise.

In addition, the shape of the transition pathway shifts appreciably under multiplicative noise. Instead of taking the straight path as under a purely deterministic regime, or the same undulating path as under additive noise, the transition pathway, manifested by the probability current, goes predominantly through the lower region of the state space. The closer the cell is around the lower region of the accessible state-space, the more likely it will be directly passed into attracting state *B*. This directional bias again arises from the coupling between the intrinsic multiplicative noise and the deterministic forces governing the system. These findings highlight the role of intrinsic multiplicative noise in the reshaping the dynamic of transitions. While the fundamental properties of the system remains intact, intrinsic multiplicative noise can lead to completely different transition dynamics by modifying the overall structure of the landscape.

## IV. DISCUSSION

Our study provides an intuitive perspective and understanding of how noise, particularly multiplicative noise, influences transition dynamics in equilibrium and non-equilibrium systems. We have here focused on exemplar systems that highlight typical behaviour and that can help us hone our intuition. Larger, and more complex systems, will exhibit similar behaviour but in a way where we cannot expect to disentangle the different contributing factors. While it is known that the interplay between noise and deterministic forces can change the structure of the landscape and give rise to qualitatuvely different features in the landscape, the influence of noise on transition dynamics, to the best of our knowledge, has not yet been studied in detail. By employing TPT in the context of Waddington’s landscape, we show that multiplicative noise can alter transition pathways by modifying the effective structure of the landscape, even when the number of attracting states (our proxy for different cell types) may be maintained.

When studying cell-state and -fate transitions, previous approaches have generally focused on the MAP, where transitions occur along minimal action paths. However, this framework can only provide information about transitions along a specific path and neglects the actual distribution of transition pathways. As we have seen, the MAP quickly becomes unrepresentative of the true dynamics: during cell differentiation small random fluctuations result in deviation from the MAP that become more severe and pronounced as the noise strength increases.

Intrinsic noise arising from the probabilistic nature of chemical reactions and gene expression is a hallmark of biological systems at the cellular, sub-cellular, and molecular level. A simplistic approach to incorporating noise is to use additive noise, an approach often favoured for its simplicity. We demonstrate that additive noise can only lead to changes in the stability of the transition state with increasing noise strength, while maintaining the underlying structure of the transition state as well as the transition pathways. However, additive noise models neglect the coupling between intrinsic noise and deterministic drift forces. More realistic forms of noise have to reflect the overall dynamics and system state. This is particularly relevant in cell differentiation where noise is known to depend on the system state: noise to be lower around the attracting states and higher near the unstable states such as saddle points.

By varying the stochastic component of a fixed deterministic system using multiplicative noise (i.e. noise that depends on the system state), we demonstrate that transition states can bifurcate and be reshaped, and so will transition pathways. Intrinsic noise not only amplifies or accelerates transitions but also, in a very concrete sense, enhances the system’s capacity to explore alternate states while preserving its fundamental structure. By examining the toggle switch model derived from the chemical Langevin equation (Eqn.(B8)), we showed that in this case, noise can alter the dominant transition pathway, even while preserving the underlying landscape structure and transition states.

Only in the simplest cases is it possible to obtain analytic results in the presence of noise. When analyic results are not amenable, particularly in high-dimensional state spaces, we must turn to numerical methods to approximate properties of interest, such as the stationary probability density function, reactive density function and mean first passage time. Here our results in two dimensions are obtained by solving Eqn. (6) using finite elements.

Our method is less effective in the small noise regime (i.e. *σ* ≪ 1). This is because the density function becomes singular at the attracting states, leading to numerical problems when computing densities directly. Furthermore, the choice of neighbourhood around attracting states significantly influences transition dynamics. Intuitively, a large choice for the basin boundary around the attracting states may encompass the area of the actual transition states, leading to different transition behaviors. In our approach, neighborhoods are defined based on specific quasi-energy levels of the corresponding deterministic systems. For practical applications involving experimental data, clustering methods or other boundary selection techniques may be necessary or preferable to accurately specify the attracting states in the system.

Our study has focused on exemplar systems for computational and illustrative purposes. However, cell differentiation typically occurs in a high-dimensional state (typically gene expression, is considered and currently the most conveniently experimentally available) space. Here transition pathways may exhibit more intricate changes under the influence of noise. We note that recent advances have introduced novel techniques for computing the statistical properties of TPT in higher dimensions [45, 46] which provide a foundation to extend our study to higher-dimensional systems.

An extension of our work to experimental data is possible in principle. The main challenge of this lies in inferring both the deterministic force and noise from experimental data. RNA velocity allows us to infer deterministic forces from scRNA-seq data, but it typically assumes additive noise [34]. To account for multiplicative noise, recent studies employ a score-matching method with neural networks, enabling the inference of deterministic force in the presence of multiplicative noise from experimental data [47, 48]. New methods of inferring gene regulatory dynamics from single-cell data that make sophisticated use of the single-cell transcriptomic data manifold offer promising routes to construct models that can be used to explore transition pathways[49].

Probability and randomness tend to defy our intuition and their effects become even more unpredicable in the context of nonlinear systems, even for the simple systems described here. At the cellular, sub-cellular, and molecular level, the effects of randomness are ubiquitously felt, including during development and cell-fate decision making processes. What we have argued here is that, as soon as biochemically relevant noise is present, the path a system takes through the relevant gene expression space can differ profoundly from what we see for the conventional deterministic case. There can be great variability among transition paths and this variability may lead to cells exploring alternative fates.

As systems become more high-dimensional, more local minima in the quasi-landscape, or more locally stable fixed points tend to arise. Understanding how developmental systems, and complex stochastic systems more generally, manage to balance stochasticity with phenotpic or overall robustness requires careful analysis within an appropriate mathematical framework. Relying on convenience – for example by using deterministic approaches or assuming purely additive noise – can obscure the underlying dynamics and be misleading. Our analysis clarifies the mathematical role of noise in shaping state transitions, providing deeper insight into cell-fate decisions within the landscape framework.

## ACKNOWLEDGMENTS

YL, AZ, and MPHS, gratefully acknowledge financial support from the Australian Research Council through a Laureate Fellowship award to MPHS. (Grant No. FL220100005).

## V. DATA AVAILABILITY

The data that support the findings of this article are openly available [50].

## Appendix A: Minimum action path

For transitions between two stable states, say *x*_initial_ and *x*_end_, from time 0 to *T*, there are many possible paths *γ*(*t*) with *γ*(0) = *x*_initial_, *γ*(*T*) = *x*_end_. In the small noise limit (i.e. *ϵ* ≪ 1), it is expected that transitions will concentrate along the path that minimizes the action functional:

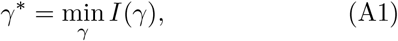

where 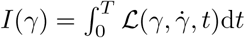 is the rate functional of path *γ* with duration *T* and 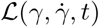 is the Lagrangian.

For systems governed by Eqn. (1), the Lagrangian can be expressed as:

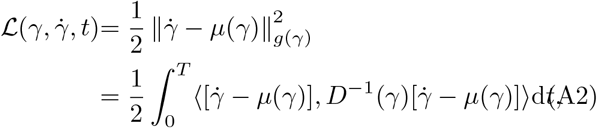

where ⟨·, ·⟩ is the Euclidean inner product and *D*^−1^ is the inverse of the diffusion matrix *D*(*x*) = *g*(*x*)*g*(*x*)^*T*^.

The probability that a cell starts at *x*_initial_ and ends at *x*_end_ at time *T* can be obtained by integrating over all possible paths between these terminal points:

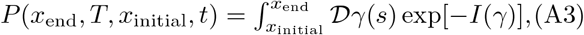

where 𝒟 *γ*(*s*) denotes the integration over all paths *γ*(*s*) with *γ*(0) = *x*_initial_ and *γ*(*T*) = *x*_end_. In the small noise regime, this probability converges to the MAP:

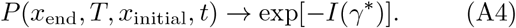

## Appendix B: Motivating problems

The landscape is generally studied using gradient systems Eqn. (2), however, biological systems function out of equilibrium at the molecular scale. In these systems, chemical energy dissipation leads to irreversible shifts between different molecular states, keeping biological systems far from thermodynamic equilibrium. Understanding these irreversible transition pathways is therefore important in understanding the mechanisms of these transition processes. The aim of this section is to give an overview of biologically relevant systems.

### 1. System in equilibrium

We start with the system that is in equilibrium. Specifically, we consider an SDE with:

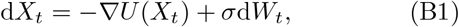

where *U* (*X*_*t*_) is some potential function and *σ* is the noise strength, which is assumed to be constant. For gradient systems, we know that the system is in equilibrium whenever additive noise is assumed [26].

The steady-state density can be expressed as the Boltzmann-Gibbs distribution:

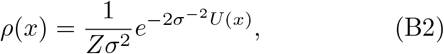

where *Z* is the normalization constant. Attracting states can be expressed as local minima in the potential function *U* (*x*). The MAP between attracting states, say *A* and *B*, will be through the saddle points separating them. Transition here is understood as the change from one state to another.

### 2. Non-equilibrium systems

We next consider more general systems that are out of equilibrium at steady state. We allow a more general SDE as given in Eqn. (1) and assume the system admits a steady density *ρ*(*x*) with

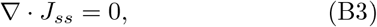

where 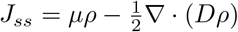 is the probability flux associated with the steady state *ρ* and *D* = *g*(*x*)*g*^*T*^ (*x*) is the diffusion matrix. When *J*_*ss*_ = 0, the system is known to be in equilibrium and the driving force under the probabilistic landscape

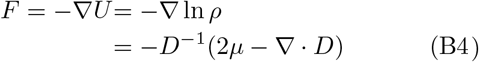

follows a perfect gradient with zero curl force:

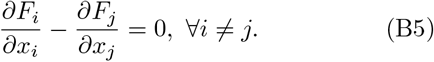

This equilibrium condition is satisfied whenever the drift term *µ* can be expressed as 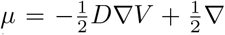 · *D* for some potential function *V* (*x*). Therefore, except in the special cases where the drift term satisfies the above conditions, the system is in non-equilibrium. This implies that the driving force *F* is composed of a gradient of the probabilistic potential force as well as a rotational curl force. Again, the attracting states are where the probabilistic landscape − ln *ρ* have local minima and the deterministic drift *µ* has stable fixed points. The transitions can be understood as the evental change from one attracting state to another after staying in one state for a long time. Note that now the MAP will be through shifted saddle points instead of the original ones due to the curl force.

### 3. Reaction networks

Finally, we consider biological systems that are described by the chemical reaction networks which are composed of many reactions. We assume there are *N* chemical species and *M* possible reactions. We denote *X*(*t*) = (*X*_1_(*t*), …, *X*_*N*_ (*t*)) as the quantities of each *i*^*th*^ species and *R*_*j*_ as the *j*^*th*^ reaction channel. When there is no noise associated with the system, the dynamics of the species evolve according to:

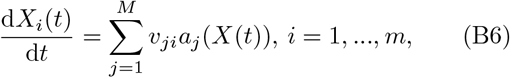

where *v*_*ij*_ ∈ ℤ^*N×M*^ gives the change in the number of *X*_*i*_ molecules under the *R*_*j*_’s reaction and the *propensity function a*_*j*_ defines the rate that *R*_*j*_’s reaction will take place.

When intrinsic noise is present, the population evolution (chemical master equation) is:

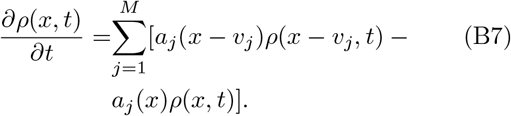

While Eqn. (B7) gives a population level description of the chemical reaction process, it is difficult to obtain the solution in general. However, one can obtain the approximated microscopic results via Ito’s form of the chemical Langevin equation (CLE):

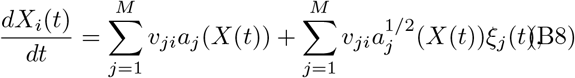

where *ξ*_*j*_(*t*) are independent Gaussian white noise terms. The underlying properties of the above systems are influenced by the intrinsic multiplicative noise where the noise *g*(*x*) is dependent on the deterministic force *µ*(*x*). We will use Eqn. (B8) to model intrinsic noise in biological reactions. We will formulate and study transitions in this system. We will also study cases where only external noise is present (i.e. the diffusion matrix *D* is constant and independent of species *X*(*t*)) and compare their effects on the transitions.

## Appendix C: Limiting behavior of reactive density in one-dimensional system with large noise strength

For the reactive density in a one-dimensional system with large noise strength

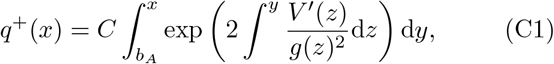

solves Eqn. (6), where *b*_*A*_ is the upper bound of the basin of attraction for state *A* and *C* ∈ ℝ ^+^ is the normalization constant that enforces *q*^+^(*x*) ∈ [0, 1], ∀*x* ∈ Ω_*AB*_.

Writing 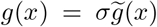, where 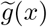 is the functional form of *g*(*x*) without scaling by *σ*, we have:

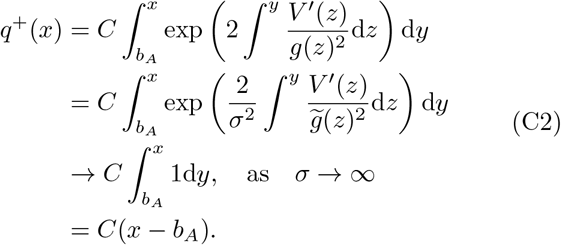

In addition, since the potential system (when 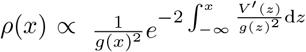) in one dimension is conservative [43], one can show that the system is time-reversible and 𝒜 = 𝒜^*^.

Therefore, the backward committor function can be expressed as:

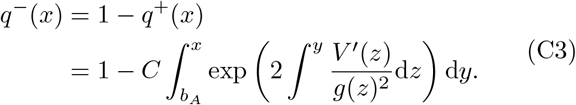

Now the reactive probability distribution under multiplicative noise can be formulated as:

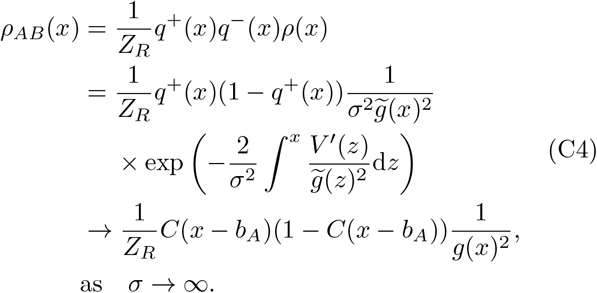

Therefore, for equilibrium systems in one dimension, as the noise strength, *σ*, increases, the reactive density will tend toward a parabolic function for additive noise. For multiplicative noise, as the noise strength increases, the reactive density will be the form of a parabolic function divided by the multiplicative noise.

## Appendix D: Finite element method for the transition property

We use the standard finite element method [51, 52] to numerically obtain the stationary distribution, committor functions and reactive density functions and transitional probability current.

The Fokker-Planck equation defines the time evolution of the probability distribution over time:

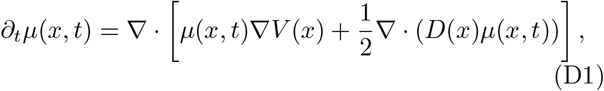

where *D*(*x*) = *g*(*X*_*t*_)^*′*^*g*(*X*_*t*_) is the diffusion matrix of the stochastic process.

By assuming ergodicity of the SDE, we have

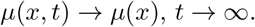

This will lead to the stationary distribution of the process defined by:

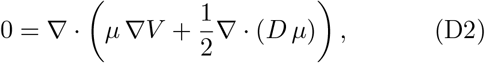

where we define *µ* = *µ*(*x*), *D* = *D*(*x*), *V* = *V* (*x*) for simplicity. Note that *µ* is constrained by ∫ _Ω_*µ* dΩ = 1. The boundary value problem for *µ* is:

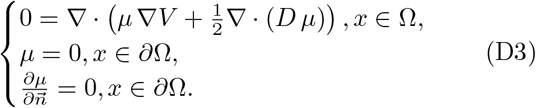

To solve this PDE with the finite element method, one needs to determine the weak formulation of this problem. Note that:

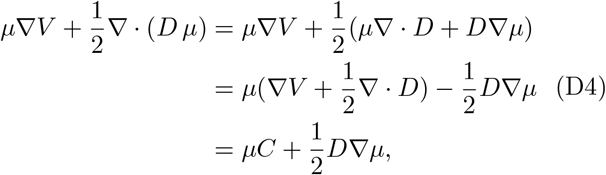

where 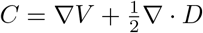. Therefore, we have:

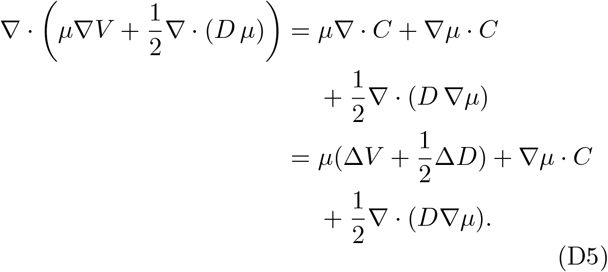

For any test function *φ* the weak formulation can be expressed as:

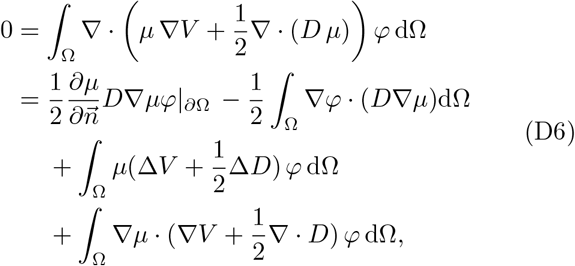

where the first term 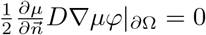 by assuming the Neumann condition at the boundary 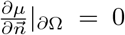. We must also constrain ∫ _Ω_*µ* dΩ = 1 to avoid a trivial 0 solution obtained by the finite element method.

We then divide the domain Ω into a finite set of elements or cells. We denote this geometric discretization grid or mesh as Ω_*h*_. Triangles are chosen as the shape for each cell, and the points where the triangles meet are referred to as nodes. We can approximate the desired value function *µ*(*x*) ≈ *µ*_*h*_(*x*) and test function *φ*(*x*) ≈*φ*_*h*_(*x*) via

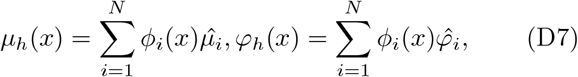

where *N* is the total number of nodes, *ϕ*_*i*_ is the basis function of each node and 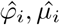 are the corresponding node values. Incorporating these approximations in the weak form we have,

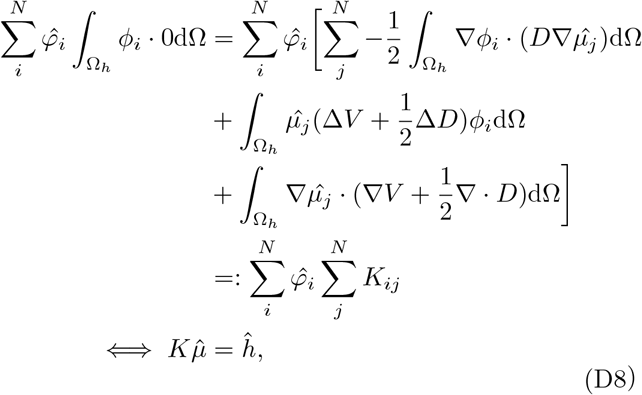

where *K* is known as the stiffness matrix and *ĥ* is the force vector with 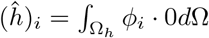.

Similarly, the boundary value problem for the forward and backward committor functions are:

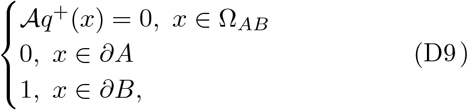

and

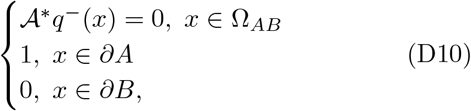

repectively. Using a similar approach as above we obtain the weak formulation for the forward committor function:

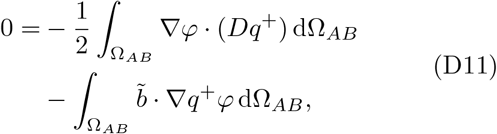

where 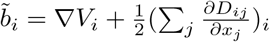.

For the backward committor function we have:

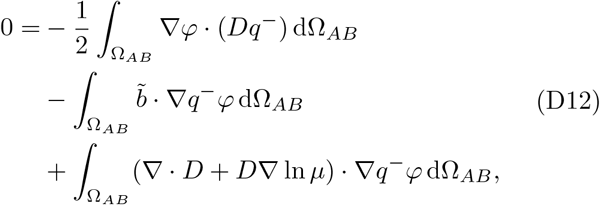

where 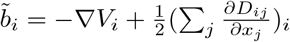.

We note that there are computational issues associated with this approach. For example, a sufficiently large choice of domain is required to constrain the probability function such that the assumption that the probability density vanishes around the boundary is valid. However, for unrealstically large domains cells may explore (stochastically and incorrectly) unrealistic regions of such unrealistic state spaces. For example, when *x, y <* − 1 in the toggle switch model, this will result in a negative value for multiplicative noise, *a*_1_(*x, y*), *a*_2_(*x, y*), and gives inaccurate results. Therefore, a careful and biologically motivated choice for the boundary is required to analyze any specific system and to obtain an accurate stationary distribution as well as reactive density.

## Appendix E: Transition pathways under higher noise strength

Here we present the reactive density of the double-well potential system under higher noise strength than in the main text. Both additive and multiplicative noise reduce the stability of the transition state, as indicated by a decrease in reactive density and an increase in flux through the transition region. However, multiplicative noise has a more profound effect: it not only destabilizes the transition state but also reshapes its structure and introduces alternative pathways towards the target state.

**FIG. E.1.**
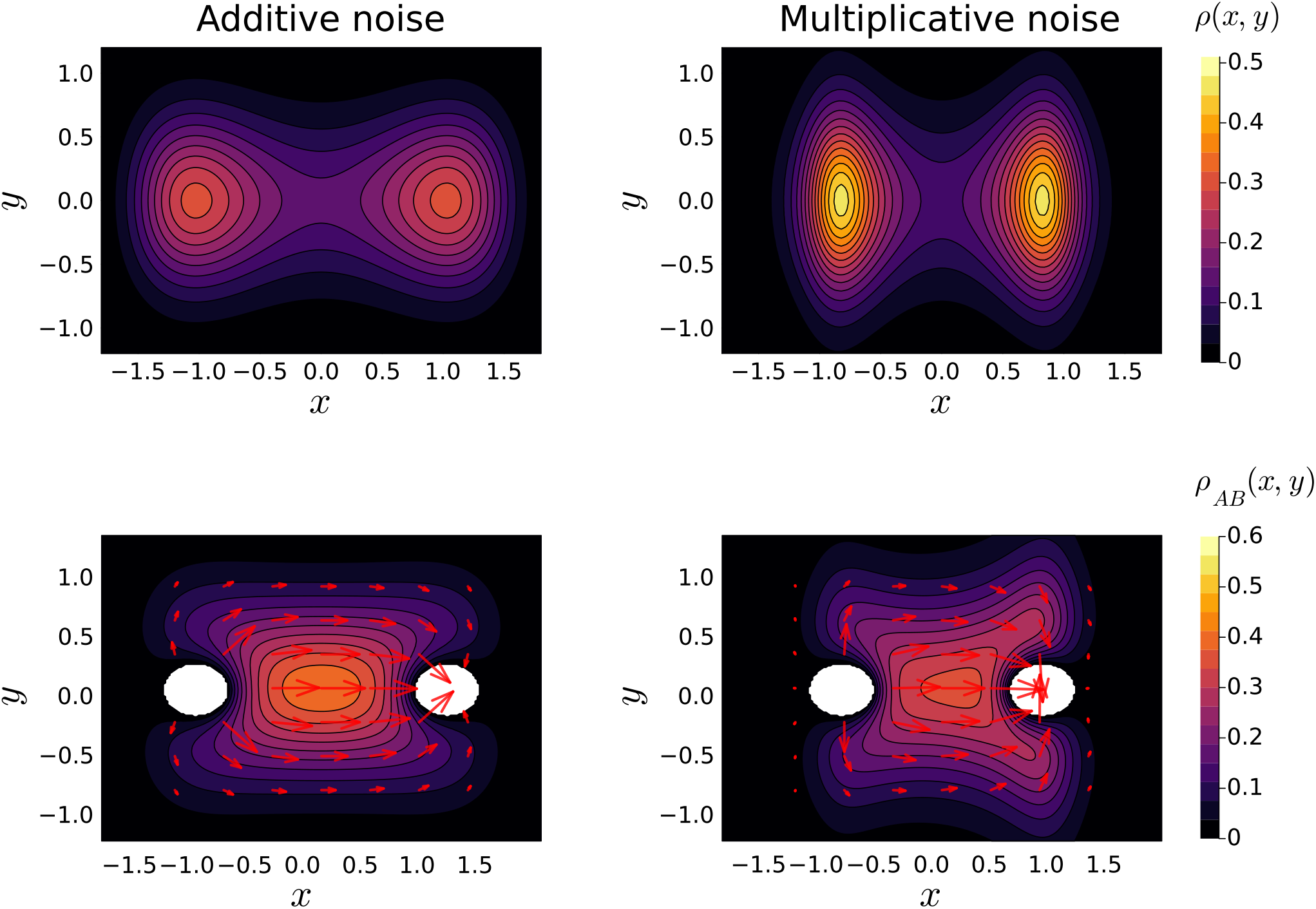
The stationary probability density function *ρ*(*x, y*) and reactive density *ρ*_*AB*_ (*x, y*) for the double-well potential system with different noise terms. (a) The stationary probability density function with additive noise described by diffusion matrix *D*_1_ on the left and multiplicative noise described by *D*_2_ on the right. (b) The reactive density *ρ*_*AB*_ (*x, y*) for with additive noise on the left and multiplicative noise on the right. The noise strength *σ* is set to be 0.8 here for both additive noise and multiplicative noise.

## References

[1] W. Zakrzewski, M. Dobrzyński, M. Szymonowicz, and Z. Rybak, Stem cells: past, present, and future, Stem Cell Research & Therapy 10 (2019).

[2] P. Rué and A. Martinez Arias, Cell dynamics and gene expression control in tissue homeostasis and development, Molecular systems biology 11, 792 (2015).

[3] L. J. Gudas and J. A. Wagner, Retinoids regulate stem cell differentiation, Journal of Cellular Physiology 226, 322 (2010).

[4] B. Alberts, Molecular biology of the cell (Garland Science, 2008).

[5] P. Haftbaradaran Esfahani and R. Knöll, Cell shape: effects on gene expression and signaling, Biophysical Reviews 12, 895 (2020).

[6] D. Burkhardt, B. Perez, J. G. Lock, S. Krishnaswamy, and C. L. Chaffer, Mapping phenotypic plasticity upon the cancer cell state landscape using manifold learning, Cancer Discovery 12, 1847 (2022).

[7] J. R. Nevins and A. Potti, Mining gene expression profiles: expression signatures as cancer phenotypes, Nature Reviews Genetics 8, 601 (2007).

[8] R. Radinsky, Modulation of tumor cell gene expression and phenotype by the organspecific metastatic environment, Cancer and Metastasis Reviews 14, 323 (1995).

[9] Y. Gong and Z. Zhang, Alternative signaling pathways: When, where and why?, FEBS Letters 579, 5265 (2005).

[10] D. Shin and K.-H. Cho, Critical transition and reversion of tumorigenesis, Experimental & Molecular Medicine 55, 692 (2023).

[11] D. Barkley, A. Rao, M. Pour, G. S. França, and I. Yanai, Cancer cell states and emergent properties of the dynamic tumor system, Genome research 31, 1719 (2021).

[12] D. Barkley, R. Moncada, M. Pour, D. A. Liberman, A. Dryg, G. Werba, W. Wang, M. Baron, A. Rao, B. Xia, G. S. França, A. Weil, D. F. Delair, C. Hajdu, A. W. Lund, I. Osman, and I. Yanai, Cancer cell states recur across tumor types and form specific interactions with the tumor microenvironment, Nature Genetics 54, 1192 (2022).

[13] R. Hernandez-Benitez, C. Wang, L. Shi, Y. Ouchi, C. Zhong, T. Hishida, H.-K. Liao, E. A. Magill, S. Memczak, R. D. Soligalla, C. Fresia, F. Hatanaka, V. Lamas, I. Guillen, S. Sahu, M. Yamamoto, Y. Shao, A. Aguirre-Vazquez, E. Nunez Delicado, P. Guillen, C. Rodriguez Esteban, J. Qu, P. Reddy, S. Horvath, G.-H. Liu, P. Magistretti, and J. C. Izpisua Belmonte, Intervention with metabolites emulating endogenous cell transitions accelerates muscle regeneration in young and aged mice, Cell Reports Medicine 5, 101449 (2024).

[14] A. Konkimalla, S. Konishi, L. Macadlo, Y. Kobayashi, Z. J. Farino, N. Miyashita, L. E. Haddad, J. M. Morowitz, C. E. Barkauskas, P. K. Agarwal, T. Souma, M. K. ElMallah, A. Tata, and P. R. Tata, Transitional cell states sculpt tissue topology during lung regeneration, Cell Stem Cell 30, 1486 (2023).

[15] Y. Mao and S. A. Wickström, Mechanical state transitions in the regulation of tissue form and function., Nat Rev Mol Cell Biol 25, 654 (2024).

[16] S. Huang, The molecular and mathematical basis of waddington’s epigenetic landscape: A framework for post-darwinian biology?, BioEssays 34, 149 (2011).

[17] J. Wang, K. Zhang, L. Xu, and E. Wang, Quantifying the waddington landscape and biological paths for development and differentiation, Proceedings of the National Academy of Sciences 108, 8257 (2011).

[18] L. Xu, K. Zhang, and J. Wang, Exploring the mechanisms of differentiation, dedifferentiation, reprogramming and transdifferentiation, PLOS ONE 9, 1 (2014).

[19] L. Xu and J. Wang, Curl flux as a dynamical origin of the bifurcations/phase transitions of nonequilibrium systems: Cell fate decision making, The Journal of Physical Chemistry B 124, 2549 (2020), pMID: 32118436, 10.1021/acs.jpcb.9b11998.

[20] K. Hayashi, H. Ohta, K. Kurimoto, S. Aramaki, and M. Saitou, Reconstitution of the mouse germ cell specification pathway in culture by pluripotent stem cells, Cell 146, 519 (2011).

[21] M. A. Coomer, L. Ham, and M. P. Stumpf, Noise distorts the epigenetic landscape and shapes cell-fate decisions, Cell Systems 10.1016/j.cels.2021.09.002 (2021).

[22] A. Guillemin and M. P. H. Stumpf, Noise and the molecular processes underlying cell fate decision-making., Physical biology 18, 011002 (2020).

[23] A. Guillemin and M. P. H. Stumpf, Non-equilibrium statistical physics, transitory epigenetic landscapes, and cell fate decision dynamics, Mathematical Biosciences and Engineering 17, 7916 (2020).

[24] L. Xu, K. Zhang, and J. Wang, Exploring the mechanisms of differentiation, dedifferentiation, reprogramming and transdifferentiation, PLOS ONE 9, 1 (2014).

[25] C. Fox, An introduction to the calculus of variations (Dover Publications, 2013).

[26] A. Bovier and F. den Hollander, Metastability (Springer, 2016).

[27] L. Cheng, X. Li, F. Li, and T. Li, Constructing the energy landscape for genetic switching system driven by intrinsic noise, PLoS One 9, e88167 (2014).

[28] C. Lv, X. Li, F. Li, and T. Li, Energy landscape reveals that the budding yeast cell cycle is a robust and adaptive multi-stage process, PLOS Computational Biology 11, e1004156 (2015).

[29] L. Ye, J. Feng, and C. Li, Controlling brain dynamics: Landscape and transition path for working memory, PLOS Computational Biology 19, 1 (2023).

[30] D. K. Wells, W. L. Kath, and A. E. Motter, Control of stochastic and induced switching in biophysical networks, Phys. Rev. X 5, 031036 (2015).

[31] W. Wang, D. Poe, Y. Yang, T. Hyatt, and J. Xing, Epithelial-to-mesenchymal transition proceeds through directional destabilization of multidimensional attractor, eLife 11, e74866 (2022).

[32] P. Zhou, S. Wang, T. Li, and Q. Nie, Dissecting transition cells from single-cell transcriptome data through multiscale stochastic dynamics, Nature Communications 12, 10.1038/s41467-021-25548-w (2021).

[33] F. Noé, C. Schütte, E. Vanden-Eijnden, L. Reich, and T. R. Weikl, Constructing the equilibrium ensemble of folding pathways from short off-equilibrium simulations, Proceedings of the National Academy of Sciences 106, 19011 (2009).

[34] X. Qiu, Y. Zhang, J. D. Martin-Rufino, C. Weng, S. Hosseinzadeh, D. Yang, A. N. Pogson, M. Y. Hein, K. Hoi (Joseph) Min, L. Wang, E. I. Grody, M. J. Shurtleff, R. Yuan, S. Xu, Y. Ma, J. M. Replogle, E. S. Lander, S. Darmanis, I. Bahar, V. G. Sankaran, J. Xing, and J. S. Weissman, Mapping transcriptomic vector fields of single cells, Cell 185, 690 (2022).

[35] W. Wang, K. Ni, D. Poe, and J. Xing, Transiently increased coordination in gene regulation during cell phenotypic transitions, PRX Life 2, 10.1103/prxlife.2.043009 (2024).

[36] R. D. Brackston, E. Lakatos, and M., Transition state characteristics during cell differentiation, PLOS Computational Biology 14, e1006405 (2018).

[37] S. Lapidus, B. Han, and J. Wang, Intrinsic noise, dissipation cost, and robustness of cellular networks: the underlying energy landscape of MAPK signal transduction, Proc. Natl. Acad. Sci. U. S. A. 105, 6039 (2008).

[38] W. E. W. Ren, and E. Vanden-Eijnden, Transition pathways in complex systems: Reaction coordinates, isocommittor surfaces, and transition tubes, Chemical Physics Letters 413, 242 (2005).

[39] W. E. and E. Vanden-Eijnden, Towards a theory of transition paths, Journal of Statistical Physics 123, 503 (2006).

[40] J. Lu and J. Nolen, Reactive trajectories and the transition path process, Probability Theory and Related Fields 161, 195 (2014).

[41] M. Ferrario, G. Ciccotti, and K. Binder, Computer Simulations in Condensed Matter Systems: From Materials to Chemical Biology Volume 1 (Springer Science+Business Media, 2006).

[42] W. E and E. Vanden-Eijnden, Transition-path theory and path-finding algorithms for the study of rare events, Annual Review of Physical Chemistry 61, 391 (2010).

[43] J. Kent, Time-reversible diffusions, Advances in Applied Probability 10, 819 (1978).

[44] F. Chen, Y. Bai, and C. Li, Estimation of nonequilibrium transition rate from gene expression data, Briefings in Bioinformatics 24, 10.1093/bib/bbad113 (2023).

[45] L. Evans, M. K. Cameron, and P. Tiwary, Computing committors via mahalanobis diffusion maps with enhanced sampling data, The Journal of Chemical Physics 157, 10.1063/5.0122990 (2022).

[46] Q. Li, B. Lin, and W. Ren, Computing committor functions for the study of rare events using deep learning, The Journal of Chemical Physics 151, 10.1063/1.5110439 (2019).

[47] V. Chardes, S. Maddu, and M. J. Shelley, Stochastic force inference via density estimation (2023), arXiv:2310.02366.

[48] A. Hyvärinen, Estimation of non-normalized statistical models by score matching, Journal of Machine Learning Research 6, 695 (2005).

[49] S. Y. Zhang and M. P. H. Stumpf, Learning cell-specific networks from dynamics and geometry of single cells, bioRxiv 10.1101/2023.01.08.523176 (2024), https://arxiv.org/abs/https://www.biorxiv.org/content/early/2024/11/13/2023.01.08.523176.full.pdf

[50] Y. Liu, yujingll/transition_pathway_under_noise: Transition pathways under noise (2025).

[51] M. Fukushima, Y. Oshima, and M. Takeda, Dirichlet Forms and Symmetric Markov Processes (De Gruyter, Berlin, New York, 2010).

[52] M. Röckner and Z.-M. Ma, Introduction to the Theory of (Non-Symmetric) Dirichlet Forms (Springer Nature, 1992).

